# Cystathionine β-synthase deficiency impairs vision in the fruit fly, *Drosophila melanogaster*

**DOI:** 10.1101/2020.03.04.975391

**Authors:** Marycruz Flores-Flores, Leonardo Moreno-García, Felipe Ángeles Castro-Martínez, Marcos Nahmad

**Affiliations:** Department of Physiology, Biophysics and Neurosciences, Center for Research and Advanced Studies, Mexico City, Mexico

**Keywords:** cystathionine β-synthase, classic homocystinuria, *Drosophila melanogaster*, phototaxis, myopia

## Abstract

**Purpose:** In humans, deficiency in Cystathionine β-Synthase (CBS) levels leads to an abnormal accumulation of homocysteine and results in classic homocystinuria, a multi-systemic disorder affecting connective tissue, muscles, the central nervous system and the eyes. However, the genetic and molecular mechanisms underlying vision problems in patients with homocystinuria are little understood.

**Materials and Methods:** The fruit fly, *Drosophila melanogaster*, is a useful experimental system to investigate the genetic basis of several human diseases, but no study to date has used Drosophila as model of homocystinuria. Here we use genetic tools to down-regulate CBS and classical behavioral assays to propose Drosophila as a model of homocystinuria to study vision problems.

**Results:** We present evidence that CBS-deficient flies show an abnormal stereotypical behavior of attraction towards a luminous source or phototaxis, consistent with severe myopia in humans. We show that this behavior cannot be fully attributed to a motor or olfactory deficiency but most likely to an impaired vision. CBS-deficient flies are overall smaller, but smaller eyes do not explain their erratic phototactic response.

**Conclusions:** We propose Drosophila as a useful model to investigate ocular manifestations underlying homocystinuria.

## Introduction

Classic homocystinuria is a metabolic disease mainly caused by inherited deficiency of Cystathionine-β-synthase (CBS), a vitamin B_6_-dependent enzyme in the transsulfuration pathway that catalyzes the flux of sulfur from methionine to cysteine ^1^. In humans, genetic variants causing low CBS expression lead to the accumulation of toxic levels of homocysteine in urine and plasma, mainly affecting skeletal, visual, and central nervous systems ^2,3^, and also poses an independent risk factor for thrombosis and vascular disease ^4,5^. One of the most common clinical manifestations of classic homocystinuria is severe myopia followed by ectopia lentis, affecting about 90% of patients with a CBS deficiency ^6,7^. Despite the high prevalence of eye-related abnormalities caused by this disease, the molecular mechanisms that relate CBS deficiency to vision problems are poorly understood. Murine models of genetic deficiency of *cbs* have been used as a model of homocystinuria ^8,9^, including its consequences in the eye. For instance, studies using *cbs*-mutant mice have reported alterations of retinal vasculature ^10^, retinal ganglion cell death ^11,12^, and visual function ^13^. However, the use of this experimental model is challenging due to a large degree of neonatal lethality ^9^.

*Drosophila melanogaster* is a useful model organism to investigate the genetic and molecular basis of development, behavior, and disease. Compared to vertebrate models, it is easier and cheaper to culture and maintain; it has a much shorter life cycle; and, offers a broad genetic toolkit to manipulate gene expression in space and time ^14^. In 2011, Kabil et al. used Drosophila to investigate the role of CBS enzyme activity on longevity and found that it is required for caloric-restricted lifespan extension ^15^. However, no study to date has reported a phenotype of CBS-deficient flies related to homocystinuria manifestations. Here, we aimed to investigate the behavior of CBS-deficient flies using a ubiquitous and eye-specific driver to express an interference RNA of *cbs* (cbs^RNAi^). Particularly, we conducted behavioral studies to ask if Drosophila vision is affected in cbs^RNAi^ mutants. We took advantage of one of the best-documented behavior in Drosophila, its response to a light stimulus or phototaxis ^16^. We assayed the phototactic response of flies expressing cbs^RNAi^ ubiquitously or in the eyes, with respect to a control group with normal CBS activity. We found significant differences in their phototactic behavior. Furthermore, we confirmed that the behavior is not due to motor impairment, suggesting that vision is affected in these mutants. Since, vision problems are a widespread manifestation in classic homocystinuria patients, we propose to establish *Drosophila melanogaster* as an animal model of homocystinuria.

## Materials and Methods

In order to downregulate CBS both, ubiquitously or in an eye-specific manner, we took advantage of the Drosophila Gal4-UAS system ^17^. We used the following fly stocks from the Bloomington Drosophila Stock Center: #36767: y, sc, v, sev;; UAS-cbs^RNAi^ (TRiP. HM-S03028), referred below as simply UAS-cbs^RNAi^. #5535: w; ey-Gal4/CyO. #25374: y, w; Act5C-Gal4/CyO. In all of our experiments, we selected non-CyO adult females resulting from the following crosses:

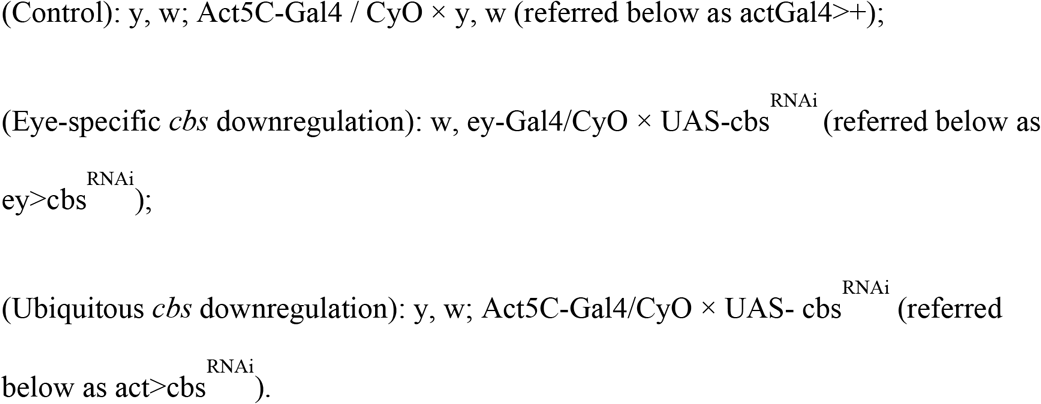

The phototactic device is a clear acrylic tube (60 cm of length × 4.5 cm of diameter). We introduced groups of 25-30 CO_2_-anesthetized flies of each genotype into one end of the tube and placed the stimulus at the other end (Figure 1A). Once the anesthesia passed in all the flies, the tube was tapped so that all the flies move to the end opposite of the stimulus and the device was immediately placed in a dark box to avoid that environmental light affect the experiment. Flies were allowed to respond to the stimulus for 10 minutes, then the box was opened and the percentage of flies located within 10 cm of the stimulus were counted (% of responsive flies; Figure 1). The experiment was repeated 5 times with the same group of flies. Each repetition was separated by a 3-minute rest time; during this time the flies were left in the device with no stimulus.

**Figure 1.**
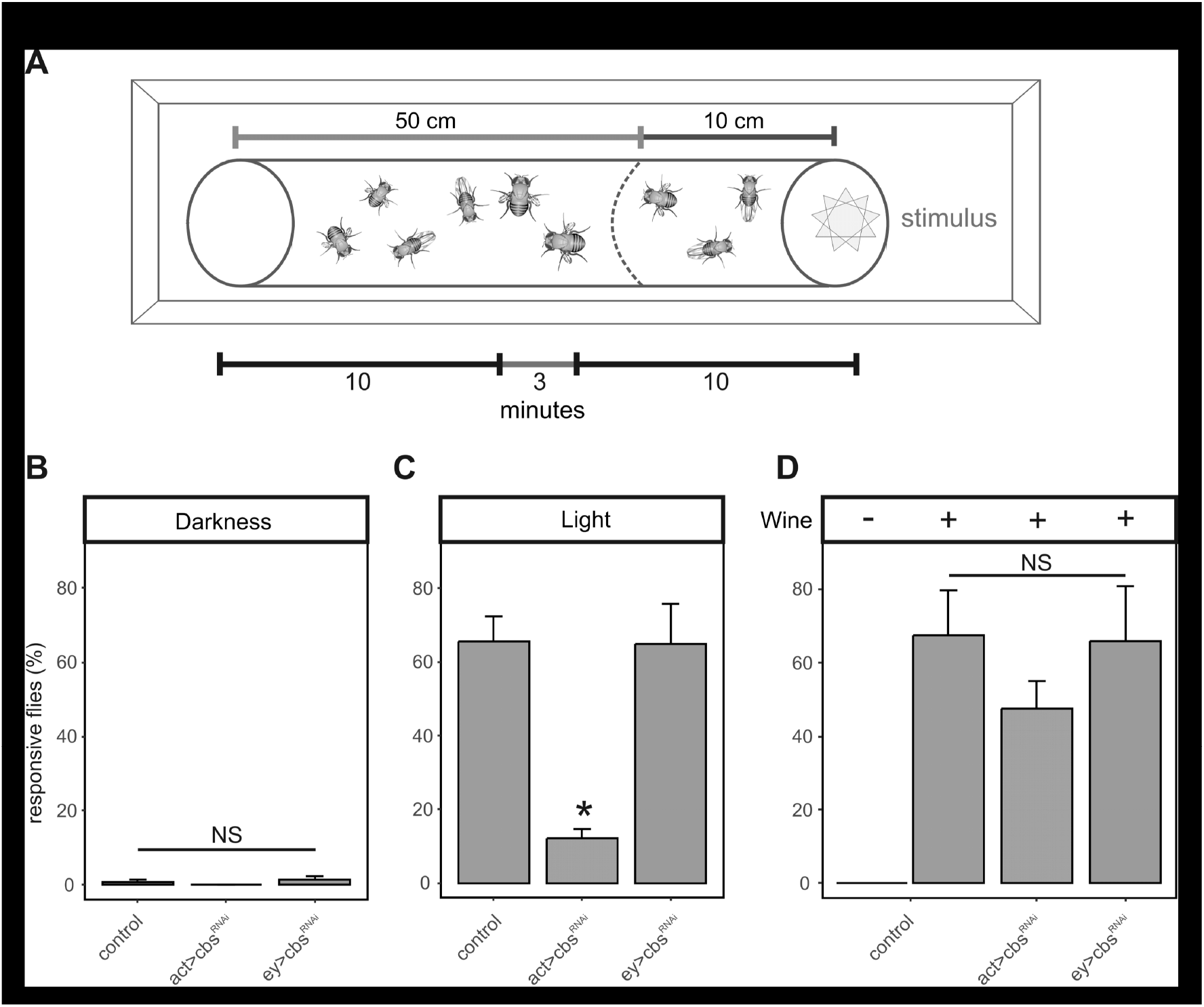
CBS-deficient flies do not exhibit the stereotypical phototaxis behavior. A. Experimental design. Diagram of the device used to evaluate the visual behavior of flies under different stimuli. Flies of each genotype are initially placed at one end of the device opposite to the location of the stimulus. Flies were allowed to respond to the stimulus for 10 minutes in a dark box to avoid environmental light and the flies located within the 10 cm mark at the stimulus end were manually counted (responsive flies). A 3-minute rest was given before starting the experiment again with the same group of flies (n=5, for each experimental group). B-D. Percentage of responding flies in the act>+ control (red bars), act>cbs^RNAi^ (green bars), ey>cbs^RNAi^ (blue bars) groups in the absence of any stimulus (B), with a white light stimulus (C), or an olfactory stimulus (D). * p<0.05, one-way ANOVA test, Tukey HSD post-hoc.

For the phototaxis experiments, the stimulus was a white LED. For the evaluation of the motor response, the LED was replaced with a cotton ball soaked in 5 mL of fresh red wine. In order to measure eyes and bodies in adults, female flies were treated overnight in a 70% ethanol solution. Both were dissected and mounted in 50% ethanol. Whole flies were placed laterally in glass coverslides. Photos in Figure 2 were taken in a Nikon ® H550S stereoscopic microscope. Measurements were taken along the axes shown in Figure 2A, B using ImageJ (https://imagej.nih.gov/ij/).

**Figure 2.**
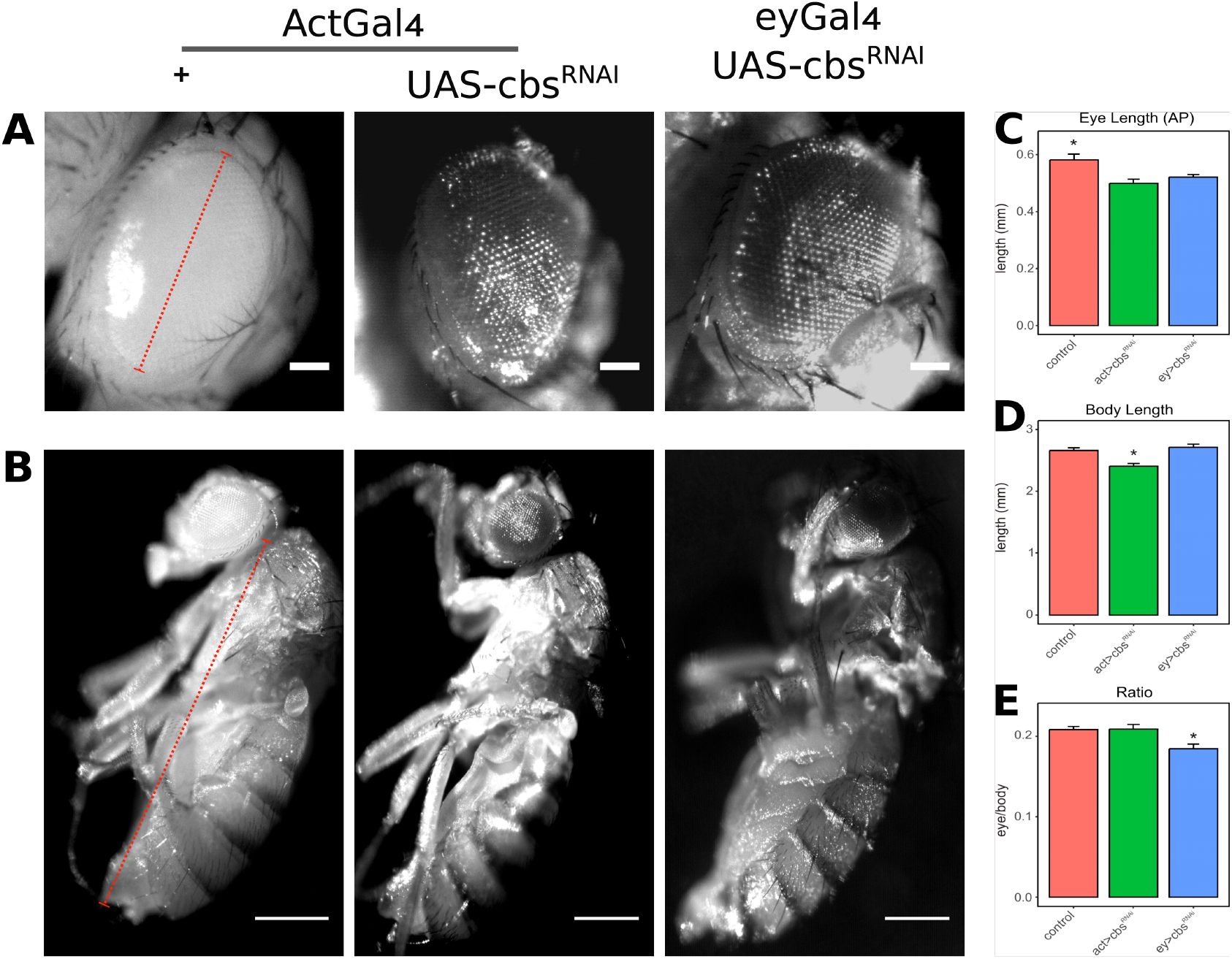
Eye size cannot explain the phototactic behaviour of CBS-deficient flies. A, B. Representative light micrographs of eyes (A, scale bar: 0.1 mm) and lateral view of the whole body (B, scale bar: 0.5 mm) of control (act>+), act>cbs^RNAi^, and ey>cbs^RNAi^ female flies. c-e. Comparison of eyes (C), body lengths (D) and the eye length to body length ratio (E) in control (red bars), act>cbs^RNAi^ (green bars), ey> cbs^RNAi^ (blue bars) female flies. The eye length was obtained by manually fitting an ellipse to the eye contour and computing the major axis of the ellipse (as indicated by the red dotted line in A). Body length was measured by computing a straight line from the head-thorax joint to the tail (as indicated by the red dotted line in B). * p<0.05, one-way ANOVA test, Tukey HSD post-hoc.

## Results and Discussion

In order to evaluate the visual response of CBS-deficient flies, we used a chamber in which flies were initially placed in one end and a light stimulus in the other end (Figure 1A). We assayed the stereotypical phototactic behavior (*i.e.*, the percentage of flies that displaced towards the light source) in three genetic conditions: control (CBS-unaffected flies; red bars), ubiquitous CBS-down-regulation (act>cbs^RNAi^; green bars), and eye-specific CBS-downregulation (ey>cbs^RNAi^; blue bars). As expected, without any stimulus, *i.e.*, when the phototactic chamber was kept in the dark, there was no significant displacement of flies to the other end of the chamber in any of the groups (Figure 1B). We then placed a light source at the end of the chamber and compared their phototactic behavior. On average, more than 60% of flies in the act>+ control group were responsive to the stimulus (Figure 1C), consistent with the classic phototactic behavior ^16^. This percentage was similar in ey>cbs^RNAi^ flies suggesting that local downregulation of CBS in eyes does not have an abnormal phototactic phenotype (Figure 1C). However, when CBS was downregulated ubiquitously (act>cbs^RNAi^) the percentage of flies that respond to the light stimulus was significantly reduced (Figure 1C), resembling vision abnormalities due to a genetic CBS deficiency in humans ^18^. In order to verify that this phenotype is specific to vision and not a motor impairment, we performed a test in which the attractant stimulus was not visual, but olfactory by replacing the light source for a cotton ball soaked in red wine, which is known to attract flies ^19^. We found that while act>cbs^RNAi^ flies appeared to respond slightly less than the control or ey>cbs^RNAi^ groups, their difference is not statistically significant (Figure 1D). We conclude that act>cbs^RNAi^ flies do not appear to have a motility or olfactory phenotype, and suggest that the impaired phototactic response is due to a defective vision.

In insects, eye size and shape are determinant factors in their visual performance ^20^. Therefore, we asked if the defective phototatic phenotype of act>cbs^RNAi^ flies could be explained by smaller eyes that would have a decrease in visual acuity ^21^. Indeed, act>cbs^RNAi^ flies were significantly and proportionally smaller than controls, as measured by whole-body and eye length (Figure 2). The reduction in CBS levels appeared to have a cell-autonomous effect on cell size and/or cell proliferation, because in ey>cbs^RNAi^ flies only eyes were significantly smaller (Figure 2C-E, compare blue bars to red and green bars). However, simply having smaller eyes did not explain a reduced phototactic response since ey>cbs^RNAi^ flies responded to a light source as well as control flies (Figure 1C). Taken together, we conclude that CBS-deficient flies have an abnormal vision that cannot be simply explained by eye size and is most likely a defect in phototransduction. Our work suggests that vision problems arising as a consequence of reduced CBS activity are evolutionary conserved from insects to humans and given the widespread genetic tools and advantages of Drosophila as a model organism, we propose its use to further investigate homocystinuria-related eye manifestations.

## Acknowledgements

This work was funded with institutional support from the Center for Research and Advanced Studies (Cinvestav). M.F-F and L.M-G were recipients of a Masters of Science scholarship from the Consejo Nacional de Ciencia y Tecnología of Mexico (CONACyT).

